## Supplemental Material for "Understanding molecular mechanisms and predicting phenotypic effects of pathogenic tubulin mutations"

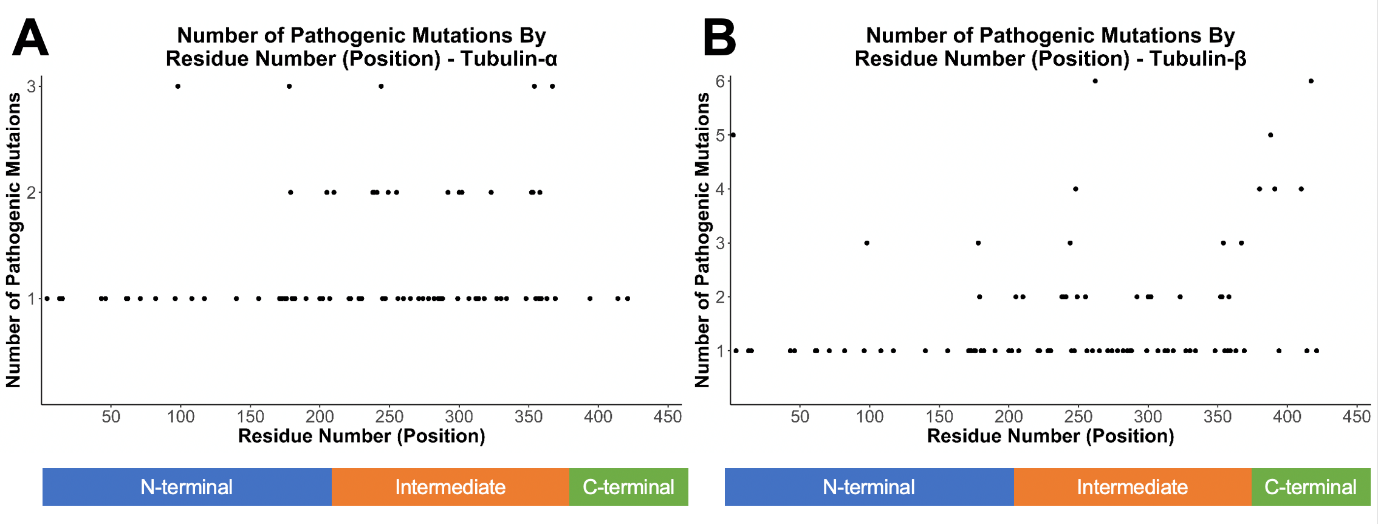

**Figure S1: Occurrence of pathogenic mutations in tubulin.**

Number of pathogenic mutations according to the residue position for tubulin-α **(A)** and β **(B)**.

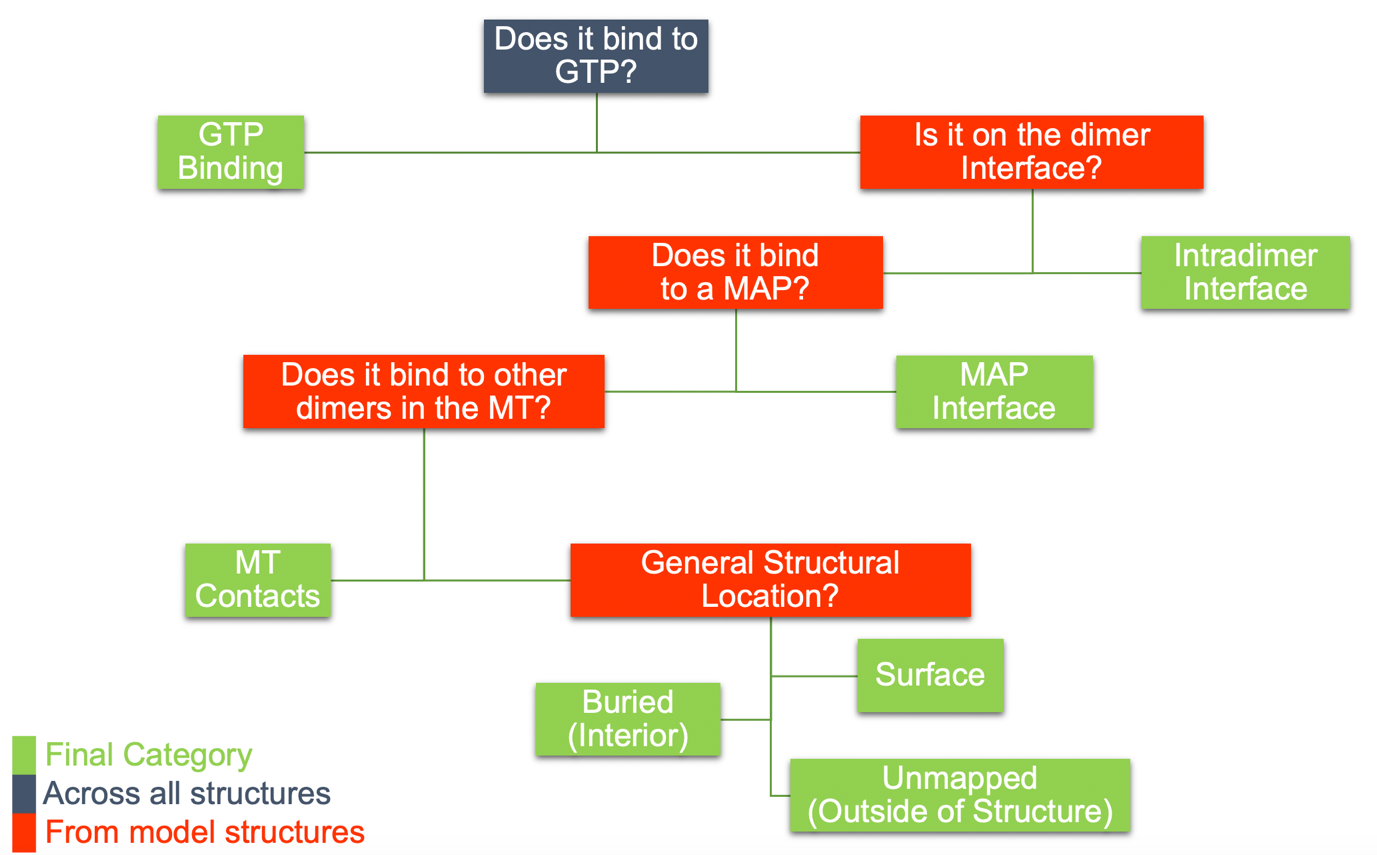

**Figure S2: Hierarchy diagram for classification of structural locations**

Schematic illustrating the hierarchy used for classification of structural locations, indicating where all structures with 70% protein sequence identity were considered, or where specific model structures were used (according to Table S2).

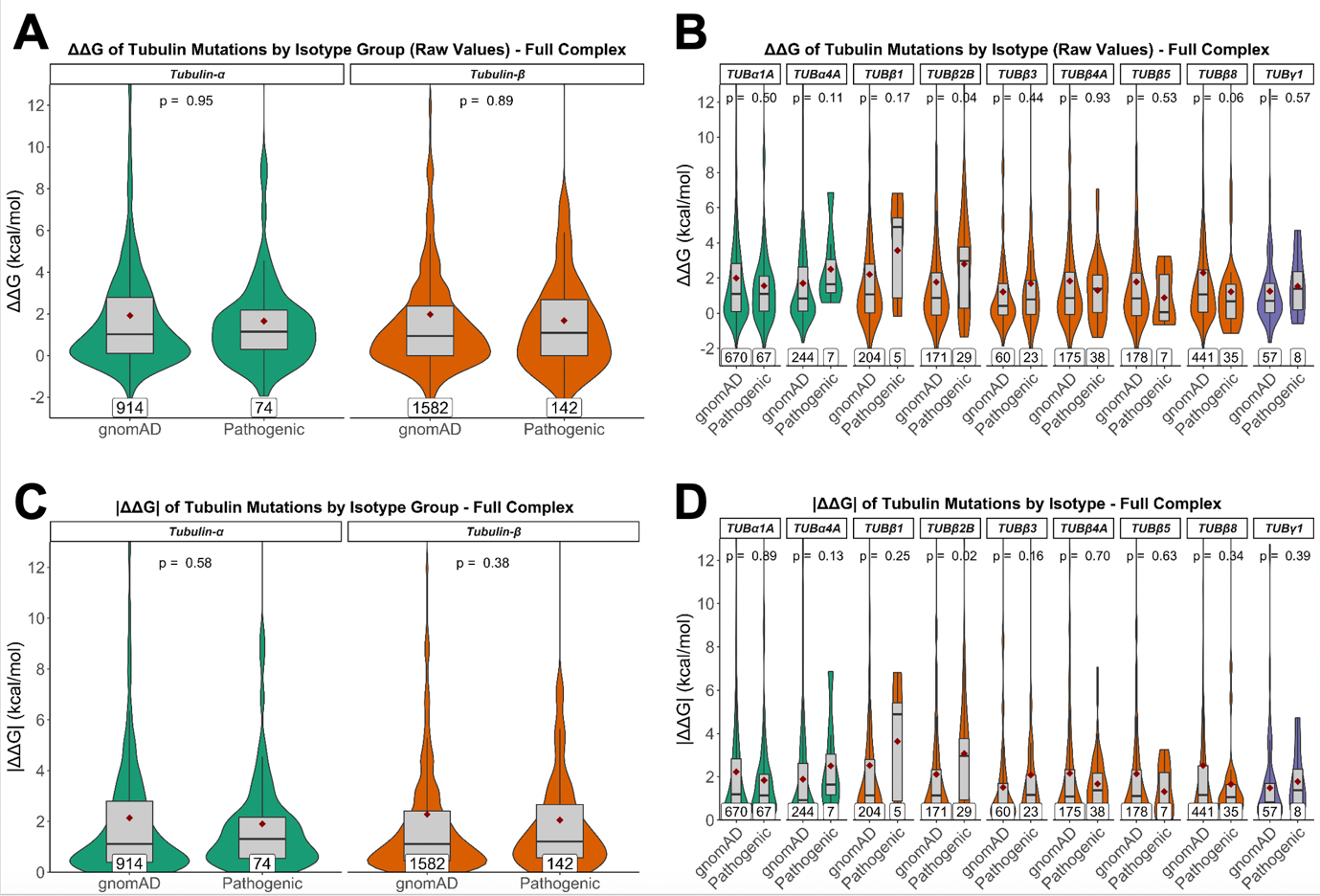

**Figure S3: Comparison of predicted changes in protein stability between pathogenic and putatively benign tubulin variants, when considering ΔΔG values calculated using full protein complex structures and absolute values**.

ΔΔG values calculating the change in free energy for folding were calculated with FoldX considering the structure of the full protien complex structures, including all intermolecular interactions. Scores are shown for tubulin-α and β families globally **(A)**, and in isotypes with at least 5 identified pathogenic mutations **(B)**. Absolute values for these scores are also shown for each family **(C)** and isotype **(D)**. Maroon diamonds indicate the mean ΔΔG score, and mutation totals for each group are also shown at the bottom. The p-values displayed were obtained via unpaired Wilcoxon tests.

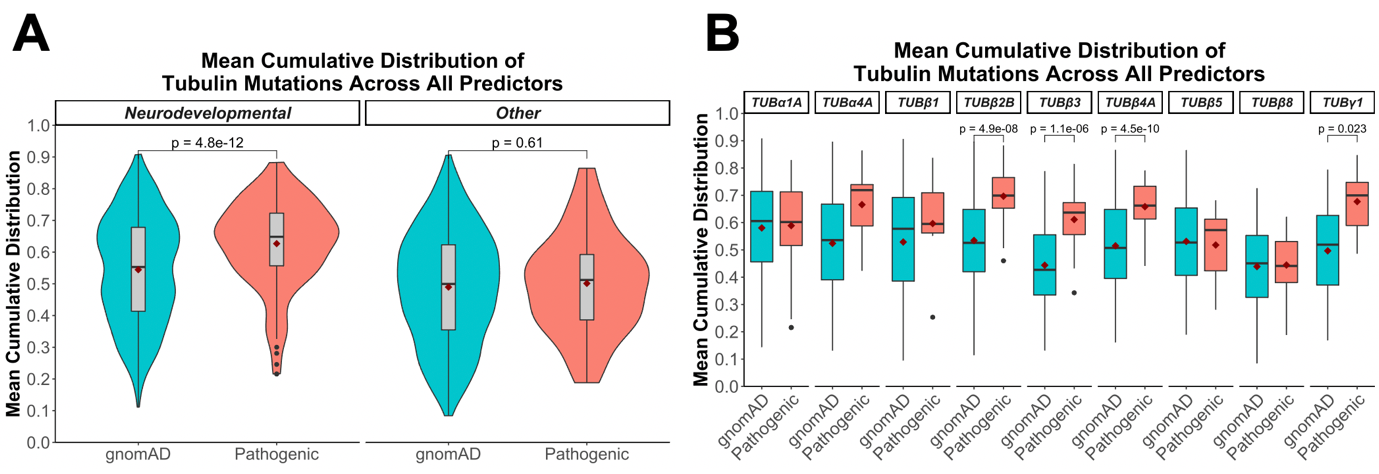

**Figure S4: Calculating the MCD scores across all predictors for each mutation reveals similar patterns**.

MCDs were calculated across all VEPs in a combined dataset which included all gnomAD and pathogenic mutations across all tubulins. The gnomAD and pathogenic MCD scores were separated and grouped by pathogenicity type **(A)**, or isotype **(B)**. The p-values stated were obtained using unpaired Wilcoxon tests, with ones not shown being above 0.05. Maroon diamonds indicate the average MCD.

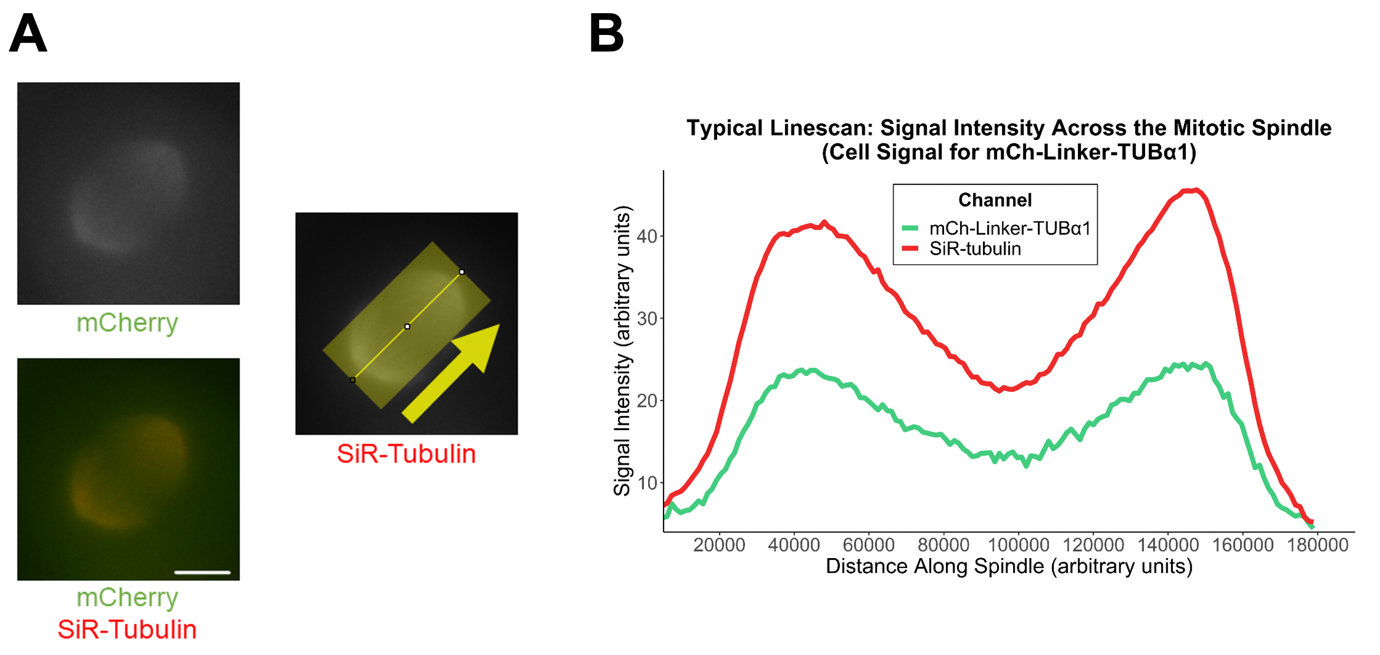

**Figure S5: Typical linescan workflow.**

**(A)** Representative live-cell images of mitotic HeLa cell after being transfected with fluorescently tagged wild type mCherry-Linker-TUBα1 (green) and incubated with SiR-tubulin (red). Using the SiR-tubulin channel, we take a wide linescan (yellow, arrow indicating direction of linescan along the x-axis) of the mitotic spindle and do the same for the fluorescent channel in each cell to determine signal intensity **(B)**. This is measured by the average background-subtracted grey value of each point across the wide line. The spearman correlation coefficient of these two traces along the linescan are then calculated on a cell-by-cell basis to account for differences of intensity.

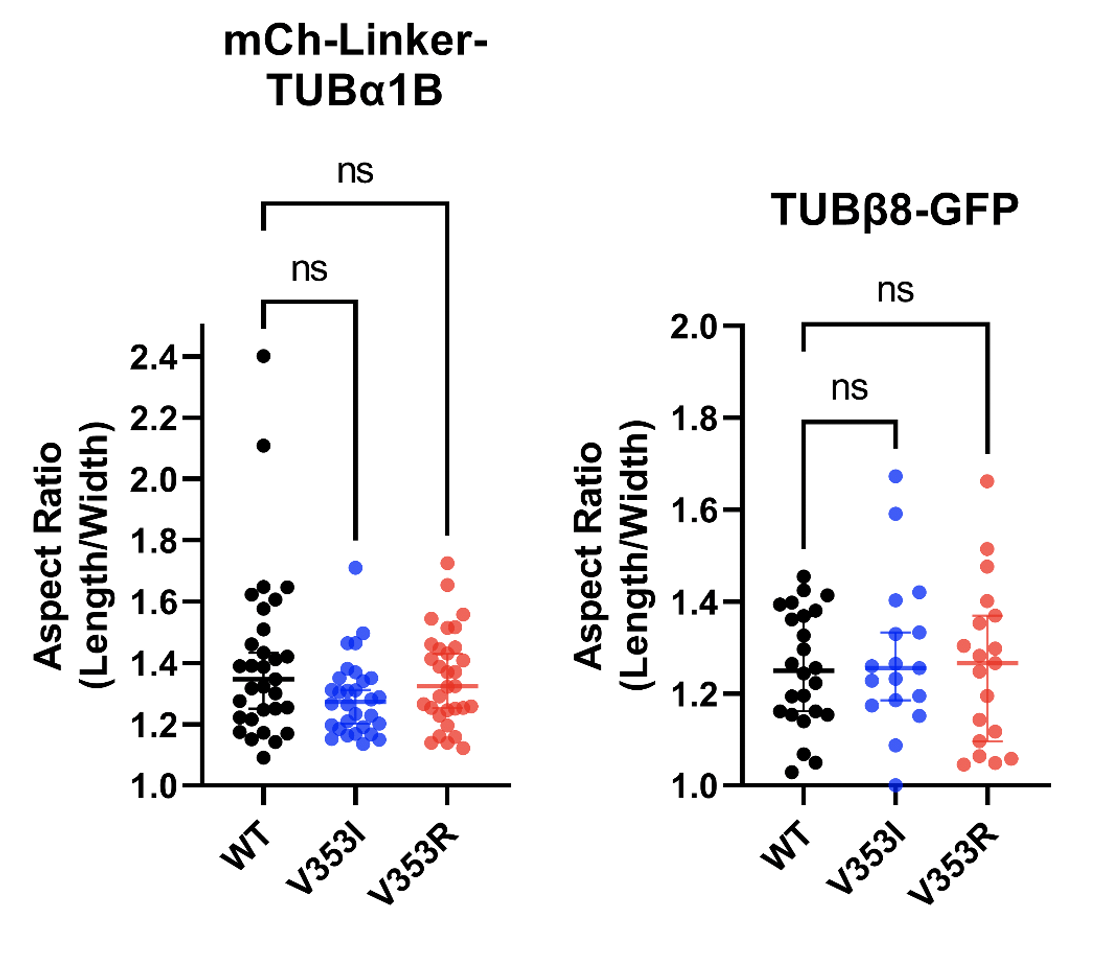

**Figure S6: Mitotic spindle width and aspect ratio of V353 mutants in TUBα1 and TUBβ8**

HeLa cells were transfected with fluorescently tagged wild type and mutant constructs and incubated with SiR-tubulin (red). With each construct, the spindle aspect ratio (length/width) was calculated for each cell. Kruskal-Wallis tests (with post-hoc Dunn) were performed, but no significant differences were found.

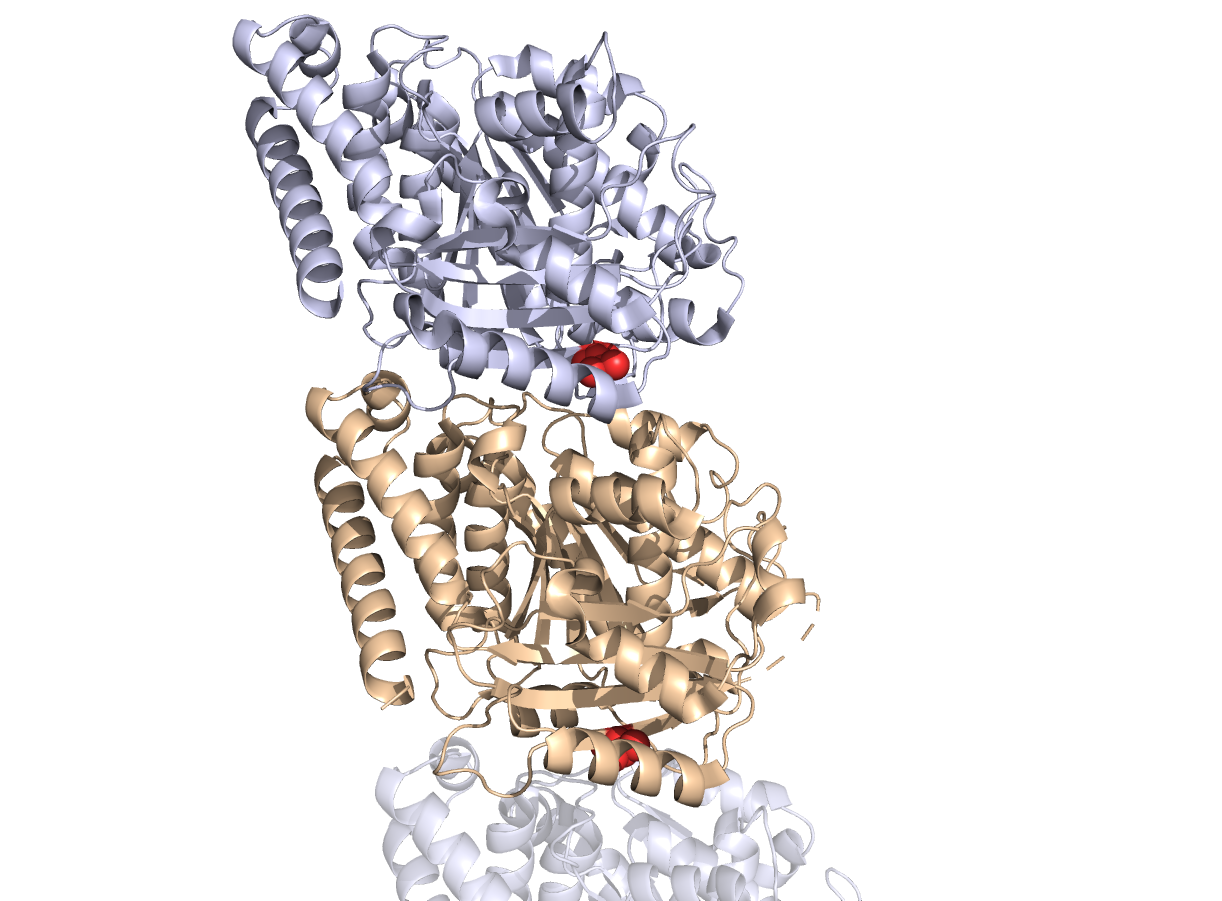

**Figure S7: Visualisation of V353 residues in tubulin-α and β.**
Tubulin-α is in wheat, while tubulin-β in bluewhite. Red spheres indicate atoms constituting the V353 residues in both proteins. Part of the next heterodimer along the microtubule protofilament is also shown in reduced transparency to show the adjacency of the V353 residue in tubulin-α to the longitudinal interdimer interface. PDB ID: 5jco

**Provided as separate files:**

**Supplemental Data 1:** Multiple sequence alignment of all tubulin-α and β isotypes considered in this study.

**Table S1:** Full description of all pathogenic mutations identified in this study (including phenotypes and references) as well as a list of all gnomAD mutations used.

**Table S2:** All model structures used for the hierarchy implemented for assigning structural locations for tubulin residues.

**Table S3:** Complete set of raw predictions obtained from all the VEPs used in this study for every tubulin mutation (gnomAD and pathogenic).

**Table S4:** DeepSequence predictions for every possible missense mutation for all tubulin isotypes considered in this study.

**Table S5:** Ranking of mean cumulative distribution (MCD) scores for all pathogenic tubulin mutations. DeepSequence scores as well as normalised ranks (preserving scale or scale-independent) also provided.

**Table S6:** All tubulin residues found to be interacting with GTP (or its analogues), MAPs, or with other tubulin subunits (interdimer or intradimer).
