## Supplemental Dataset 1 for "Understanding molecular mechanisms and predicting phenotypic effects of pathogenic tubulin mutations"

CLUSTAL O(1.2.4) Multiple Sequence Alignment – All Tubulin-α Amino Acid Sequences

```
TBA1A_HUMAN/1-451      MRECISIHVGQAGVQIGNACWELYCLEHGIQPDGQMPSDKTIGGGDDSFNTFFSETGAGK   60
TBA1B_HUMAN/1-451      MRECISIHVGQAGVQIGNACWELYCLEHGIQPDGQMPSDKTIGGGDDSFNTFFSETGAGK   60
TBA1C_HUMAN/1-449      MRECISIHVGQAGVQIGNACWELYCLEHGIQPDGQMPSDKTIGGGDDSFNTFFSETGAGK   60
TBA3D_HUMAN/1-450      MRECISIHVGQAGVQIGNACWELYCLEHGIQPDGQMPSDKTIGGGDDSFNTFFSETGAGK   60
TBA3E_HUMAN/1-450      MRECISIHVGQAGVQIGNACWELYCLEHGIQPDGQMPSDKTIGGGDDSFNTFFSETGAGK   60
TBA4A_HUMAN/1-448      MRECISVHVGQAGVQMGNACWELYCLEHGIQPDGQMPSDKTIGGGDDSFTTFFCETGAGK   60
TBA8_HUMAN/1-449       MRECISVHVGQAGVQIGNACWELFCLEHGIQADGTFDAQASKINDDDSFTTFFSETGNGK   60
                       ******:********:*******:******* ** : :: :  ..****.***.*** **

TBA1A_HUMAN/1-451      HVPRAVFVDLEPTVIDEVRTGTYRQLFHPEQLITGKEDAANNYARGHYTIGKEIIDLVLD   120
TBA1B_HUMAN/1-451      HVPRAVFVDLEPTVIDEVRTGTYRQLFHPEQLITGKEDAANNYARGHYTIGKEIIDLVLD   120
TBA1C_HUMAN/1-449      HVPRAVFVDLEPTVIDEVRTGTYRQLFHPEQLITGKEDAANNYARGHYTIGKEIIDLVLD   120
TBA3D_HUMAN/1-450      HVPRAVFVDLEPTVVDEVRTGTYRQLFHPEQLITGKEDAANNYARGHYTIGKEIVDLVLD   120
TBA3E_HUMAN/1-450      HVPRAVFVDLEPTVVDEVRTGTYRQLFHPEQLITGKEDAASNYARGHYTIGKEIVDLVLD   120
TBA4A_HUMAN/1-448      HVPRAVFVDLEPTVIDEIRNGPYRQLFHPEQLITGKEDAANNYARGHYTIGKEIIDPVLD   120
TBA8_HUMAN/1-449       HVPRAVMIDLEPTVVDEVRAGTYRQLFHPEQLITGKEDAANNYARGHYTVGKESIDLVLD   120
                       ******::******:**:* * ******************.********:*** :* ***

TBA1A_HUMAN/1-451      RIRKLADQCTGLQGFLVFHSFGGGTGSGFTSLLMERLSVDYGKKSKLEFSIYPAPQVSTA   180
TBA1B_HUMAN/1-451      RIRKLADQCTGLQGFLVFHSFGGGTGSGFTSLLMERLSVDYGKKSKLEFSIYPAPQVSTA   180
TBA1C_HUMAN/1-449      RIRKLADQCTGLQGFLVFHSFGGGTGSGFTSLLMERLSVDYGKKSKLEFSIYPAPQVSTA   180
TBA3D_HUMAN/1-450      RIRKLADLCTGLQGFLIFHSFGGGTGSGFASLLMERLSVDYGKKSKLEFAIYPAPQVSTA   180
TBA3E_HUMAN/1-450      RIRKLADLCTGLQGFLIFHSFGGGTGSGFASLLMERLSVDYSKKSKLEFAIYPAPQVSTA   180
TBA4A_HUMAN/1-448      RIRKLSDQCTGLQGFLVFHSFGGGTGSGFTSLLMERLSVDYGKKSKLEFSIYPAPQVSTA   180
TBA8_HUMAN/1-449       RIRKLTDACSGLQGFLIFHSFGGGTGSGFTSLLMERLSLDYGKKSKLEFAIYPAPQVSTA   180
                       *****:* *:******:************:********:**.*******:**********

TBA1A_HUMAN/1-451      VVEPYNSILTTHTTLEHSDCAFMVDNEAIYDICRRNLDIERPTYTNLNRLIGQIVSSITA   240
TBA1B_HUMAN/1-451      VVEPYNSILTTHTTLEHSDCAFMVDNEAIYDICRRNLDIERPTYTNLNRLISQIVSSITA   240
TBA1C_HUMAN/1-449      VVEPYNSILTTHTTLEHSDCAFMVDNEAIYDICRRNLDIERPTYTNLNRLISQIVSSITA   240
TBA3D_HUMAN/1-450      VVEPYNSILTTHTTLEHSDCAFMVDNEAIYDICRRNLDIERPTYTNLNRLIGQIVSSITA   240
TBA3E_HUMAN/1-450      VVEPYNSILTTHTTLEHSDCAFMVDNEAIYDICRRNLDIERPTYTNLNRLIGQIVSSITA   240
TBA4A_HUMAN/1-448      VVEPYNSILTTHTTLEHSDCAFMVDNEAIYDICRRNLDIERPTYTNLNRLISQIVSSITA   240
TBA8_HUMAN/1-449       VVEPYNSILTTHTTLEHSDCAFMVDNEAIYDICRRNLDIERPTYTNLNRLISQIVSSITA   240
                       ***************************************************.********

TBA1A_HUMAN/1-451      SLRFDGALNVDLTEFQTNLVPYPRIHFPLATYAPVISAEKAYHEQLSVAEITNACFEPAN   300
TBA1B_HUMAN/1-451      SLRFDGALNVDLTEFQTNLVPYPRIHFPLATYAPVISAEKAYHEQLSVAEITNACFEPAN   300
TBA1C_HUMAN/1-449      SLRFDGALNVDLTEFQTNLVPYPRIHFPLATYAPVISAEKAYHEQLTVAEITNACFEPAN   300
TBA3D_HUMAN/1-450      SLRFDGALNVDLTEFQTNLVPYPRIHFPLATYAPVISAEKAYHEQLSVAEITNACFEPAN   300
TBA3E_HUMAN/1-450      SLRFDGALNVDLTEFQTNLVPYPRIHFPLATYAPVISAEKAYHEQLSVAEITNACFEPAN   300
TBA4A_HUMAN/1-448      SLRFDGALNVDLTEFQTNLVPYPRIHFPLATYAPVISAEKAYHEQLSVAEITNACFEPAN   300
TBA8_HUMAN/1-449       SLRFDGALNVDLTEFQTNLVPYPRIHFPLVTYAPIISAEKAYHEQLSVAEITSSCFEPNS   300
                       *****************************.****:***********:*****.:**** .

TBA1A_HUMAN/1-451      QMVKCDPRHGKYMACCLLYRGDVVPKDVNAAIATIKTKRTIQFVDWCPTGFKVGINYQPP   360
TBA1B_HUMAN/1-451      QMVKCDPRHGKYMACCLLYRGDVVPKDVNAAIATIKTKRSIQFVDWCPTGFKVGINYQPP   360
TBA1C_HUMAN/1-449      QMVKCDPRHGKYMACCLLYRGDVVPKDVNAAIATIKTKRTIQFVDWCPTGFKVGINYQPP   360
TBA3D_HUMAN/1-450      QMVKCDPRHGKYMACCMLYRGDVVPKDVNAAIATIKTKRTIQFVDWCPTGFKVGINYQPP   360
TBA3E_HUMAN/1-450      QMVKCDPRHGKYMACCMLYRGDVVPKDVNAAIATIKTKRTIQFVDWCPTGFKVGINYQPP   360
TBA4A_HUMAN/1-448      QMVKCDPRHGKYMACCLLYRGDVVPKDVNAAIAAIKTKRSIQFVDWCPTGFKVGINYQPP   360
TBA8_HUMAN/1-449       QMVKCDPRHGKYMACCMLYRGDVVPKDVNVAIAAIKTKRTIQFVDWCPTGFKVGINYQPP   360
                       ****************:************.***:*****:********************


TBA1A_HUMAN/1-451      TVVPGGDLAKVQRAVCMLSNTTAIAEAWARLDHKFDLMYAKRAFVHWYVGEGMEEGEFSE   420
TBA1B_HUMAN/1-451      TVVPGGDLAKVQRAVCMLSNTTAIAEAWARLDHKFDLMYAKRAFVHWYVGEGMEEGEFSE   420
TBA1C_HUMAN/1-449      TVVPGGDLAKVQRAVCMLSNTTAVAEAWARLDHKFDLMYAKRAFVHWYVGEGMEEGEFSE   420
TBA3D_HUMAN/1-450      TVVPGGDLAKVQRAVCMLSNTTAIAEAWARLDHKFDLMYAKRAFVHWYVGEGMEEGEFSE   420
TBA3E_HUMAN/1-450      TVVPGGDLAKVQRAVCMLSNTTAIAEAWARLVHKFDLMYAKWAFVHWYVGEGMEEGEFSE   420
TBA4A_HUMAN/1-448      TVVPGGDLAKVQRAVCMLSNTTAIAEAWARLDHKFDLMYAKRAFVHWYVGEGMEEGEFSE   420
TBA8_HUMAN/1-449       TVVPGGDLAKVQRAVCMLSNTTAIAEAWARLDHKFDLMYAKRAFVHWYVGEGMEEGEFSE   420
                       ***********************:******* ********* ******************
```

```
TBA1A_HUMAN/1-451      AREDMAALEKDYEEVGVDSVEGEGEEEGEEY 451
TBA1B_HUMAN/1-451      AREDMAALEKDYEEVGVDSVEGEGEEEGEEY 451
TBA1C_HUMAN/1-449      AREDMAALEKDYEEVGADSADGEDEGEEY-- 449
TBA3D_HUMAN/1-450      AREDLAALEKDYEEVGVDSVEAEAEEGEEY- 450
TBA3E_HUMAN/1-450      AREDLAALEKDCEEVGVDSVEAEAEEGEAY- 450
TBA4A_HUMAN/1-448      AREDMAALEKDYEEVGIDSYEDEDEGEE--- 448
TBA8_HUMAN/1-449       AREDLAALEKDYEEVGTDSFEEENEGEEF-- 449
                       ****:****** **** ** : * *
```

CLUSTAL O(1.2.4) Multiple Sequence Alignment – All Tubulin-β Amino Acid Sequences

```
TBB1_HUMAN/1-451      MREIVHIQIGQCGNQIGAKFWEMIGEEHGIDLAGSDRGASALQLERISVYYNEAYGRKYV   60
TBB2A_HUMAN/1-445     MREIVHIQAGQCGNQIGAKFWEVISDEHGIDPTGSYHGDSDLQLERINVYYNEAAGNKYV   60
TBB2B_HUMAN/1-445     MREIVHIQAGQCGNQIGAKFWEVISDEHGIDPTGSYHGDSDLQLERINVYYNEATGNKYV   60
TBB4A_HUMAN/1-444     MREIVHLQAGQCGNQIGAKFWEVISDEHGIDPTGTYHGDSDLQLERINVYYNEATGGNYV   60
TBB3_HUMAN/1-450      MREIVHIQAGQCGNQIGAKFWEVISDEHGIDPSGNYVGDSDLQLERISVYYNEASSHKYV   60
TBB4B_HUMAN/1-445     MREIVHLQAGQCGNQIGAKFWEVISDEHGIDPTGTYHGDSDLQLERINVYYNEATGGKYV   60
TBB5_HUMAN/1-444      MREIVHIQAGQCGNQIGAKFWEVISDEHGIDPTGTYHGDSDLQLDRISVYYNEATGGKYV   60
TBB8_HUMAN/1-444      MREIVLTQIGQCGNQIGAKFWEVISDEHAIDSAGTYHGDSHLQLERINVYYNEASGGRYV   60
                      *****  * *************:*.:**.** :*.  * * ***:**.****** . .**

TBB1_HUMAN/1-451      PRAVLVDLEPGTMDSIRSSKLGALFQPDSFVHGNSGAGNNWAKGHYTEGAELIENVLEVV   120
TBB2A_HUMAN/1-445     PRAILVDLEPGTMDSVRSGPFGQIFRPDNFVFGQSGAGNNWAKGHYTEGAELVDSVLDVV   120
TBB2B_HUMAN/1-445     PRAILVDLEPGTMDSVRSGPFGQIFRPDNFVFGQSGAGNNWAKGHYTEGAELVDSVLDVV   120
TBB4A_HUMAN/1-444     PRAVLVDLEPGTMDSVRSGPFGQIFRPDNFVFGQSGAGNNWAKGHYTEGAELVDAVLDVV   120
TBB3_HUMAN/1-450      PRAILVDLEPGTMDSVRSGAFGHLFRPDNFIFGQSGAGNNWAKGHYTEGAELVDSVLDVV   120
TBB4B_HUMAN/1-445     PRAVLVDLEPGTMDSVRSGPFGQIFRPDNFVFGQSGAGNNWAKGHYTEGAELVDSVLDVV   120
TBB5_HUMAN/1-444      PRAILVDLEPGTMDSVRSGPFGQIFRPDNFVFGQSGAGNNWAKGHYTEGAELVDSVLDVV   120
TBB8_HUMAN/1-444      PRAVLVDLEPGTMDSVRSGPFGQVFRPDNFIFGQCGAGNNWAKGHYTEGAELMESVMDVV   120
                      ***:***********:**. :* :*:**.*:.*:.*****************:: *::**

TBB1_HUMAN/1-451      RHESESCDCLQGFQIVHSLGGGTGSGMGTLLMNKIREEYPDRIMNSFSVMPSPKVSDTVV   180
TBB2A_HUMAN/1-445     RKESESCDCLQGFQLTHSLGGGTGSGMGTLLISKIREEYPDRIMNTFSVMPSPKVSDTVV   180
TBB2B_HUMAN/1-445     RKESESCDCLQGFQLTHSLGGGTGSGMGTLLISKIREEYPDRIMNTFSVMPSPKVSDTVV   180
TBB4A_HUMAN/1-444     RKEAESCDCLQGFQLTHSLGGGTGSGMGTLLISKIREEFPDRIMNTFSVVPSPKVSDTVV   180
TBB3_HUMAN/1-450      RKECENCDCLQGFQLTHSLGGGTGSGMGTLLISKVREEYPDRIMNTFSVVPSPKVSDTVV   180
TBB4B_HUMAN/1-445     RKEAESCDCLQGFQLTHSLGGGTGSGMGTLLISKIREEYPDRIMNTFSVVPSPKVSDTVV   180
TBB5_HUMAN/1-444      RKEAESCDCLQGFQLTHSLGGGTGSGMGTLLISKIREEYPDRIMNTFSVVPSPKVSDTVV   180
TBB8_HUMAN/1-444      RKEAESCDCLQGFQLTHSLGGGTGSGMGTLLLSKIREEYPDRIINTFSILPSPKVSDTVV   180
                      *:*.*.********:.***************:.*:***:****:*:**::**********

TBB1_HUMAN/1-451      EPYNAVLSIHQLIENADACFCIDNEALYDICFRTLKLTTPTYGDLNHLVSLTMSGITTSL   240
TBB2A_HUMAN/1-445     EPYNATLSVHQLVENTDETYSIDNEALYDICFRTLKLTTPTYGDLNHLVSATMSGVTTCL   240
TBB2B_HUMAN/1-445     EPYNATLSVHQLVENTDETYCIDNEALYDICFRTLKLTTPTYGDLNHLVSATMSGVTTCL   240
TBB4A_HUMAN/1-444     EPYNATLSVHQLVENTDETYCIDNEALYDICFRTLKLTTPTYGDLNHLVSATMSGVTTCL   240
TBB3_HUMAN/1-450      EPYNATLSIHQLVENTDETYCIDNEALYDICFRTLKLATPTYGDLNHLVSATMSGVTTSL   240
TBB4B_HUMAN/1-445     EPYNATLSVHQLVENTDETYCIDNEALYDICFRTLKLTTPTYGDLNHLVSATMSGVTTCL   240
TBB5_HUMAN/1-444      EPYNATLSVHQLVENTDETYCIDNEALYDICFRTLKLTTPTYGDLNHLVSATMSGVTTCL   240
TBB8_HUMAN/1-444      EPYNATLSVHQLIENADETFCIDNEALYDICSKTLKLPTPTYGDLNHLVSATMSGVTTCL   240
                      *****.**:***:**:*  :.********** :**** ************ ****:**.*

TBB1_HUMAN/1-451      RFPGQLNADLRKLAVNMVPFPRLHFFMPGFAPLTAQGSQQYRALSVAELTQQMFDARNTM   300
TBB2A_HUMAN/1-445     RFPGQLNADLRKLAVNMVPFPRLHFFMPGFAPLTSRGSQQYRALTVPELTQQMFDSKNMM   300
TBB2B_HUMAN/1-445     RFPGQLNADLRKLAVNMVPFPRLHFFMPGFAPLTSRGSQQYRALTVPELTQQMFDSKNMM   300
TBB4A_HUMAN/1-444     RFPGQLNADLRKLAVNMVPFPRLHFFMPGFAPLTSRGSQQYRALTVPELTQQMFDAKNMM   300
TBB3_HUMAN/1-450      RFPGQLNADLRKLAVNMVPFPRLHFFMPGFAPLTARGSQQYRALTVPELTQQMFDAKNMM   300
TBB4B_HUMAN/1-445     RFPGQLNADLRKLAVNMVPFPRLHFFMPGFAPLTSRGSQQYRALTVPELTQQMFDAKNMM   300
TBB5_HUMAN/1-444      RFPGQLNADLRKLAVNMVPFPRLHFFMPGFAPLTSRGSQQYRALTVPELTQQVFDAKNMM   300
TBB8_HUMAN/1-444      RFPGQLNADLRKLAVNMVPFPRLHFFMPGFAPLTSRGSQQYRALTVAELTQQMFDAKNMM   300
                      **********************************::********:* *****:**::* *

TBB1_HUMAN/1-451      AACDLRRGRYLTVACIFRGKMSTKEVDQQLLSVQTRNSSCFVEWIPNNVKVAVCDIPPRG   360
TBB2A_HUMAN/1-445     AACDPRHGRYLTVAAIFRGRMSMKEVDEQMLNVQNKNSSYFVEWIPNNVKTAVCDIPPRG   360
TBB2B_HUMAN/1-445     AACDPRHGRYLTVAAIFRGRMSMKEVDEQMLNVQNKNSSYFVEWIPNNVKTAVCDIPPRG   360
TBB4A_HUMAN/1-444     AACDPRHGRYLTVAAVFRGRMSMKEVDEQMLSVQSKNSSYFVEWIPNNVKTAVCDIPPRG   360
TBB3_HUMAN/1-450      AACDPRHGRYLTVATVFRGRMSMKEVDEQMLAIQSKNSSYFVEWIPNNVKVAVCDIPPRG   360
TBB4B_HUMAN/1-445     AACDPRHGRYLTVAAVFRGRMSMKEVDEQMLNVQNKNSSYFVEWIPNNVKTAVCDIPPRG   360
TBB5_HUMAN/1-444      AACDPRHGRYLTVAAVFRGRMSMKEVDEQMLNVQNKNSSYFVEWIPNNVKTAVCDIPPRG   360
TBB8_HUMAN/1-444      AACDPRHGRYLTAAAIFRGRMPMREVDEQMFNIQDKNSSYFADWLPNNVKTAVCDIPPRG   360
                      **** *:*****.* :***:*  :***:*:: :* :*** *.:*:*****.*********

TBB1_HUMAN/1-451      LSMAATFIGNNTAIQEIFNRVSEHFSAMFKRKAFVHWYTSEGMDINEFGEAENNIHDLVS   420
TBB2A_HUMAN/1-445     LKMSATFIGNSTAIQELFKRISEQFTAMFRRKAFLHWYTGEGMDEMEFTEAESNMNDLVS   420
TBB2B_HUMAN/1-445     LKMSATFIGNSTAIQELFKRISEQFTAMFRRKAFLHWYTGEGMDEMEFTEAESNMNDLVS   420
TBB4A_HUMAN/1-444     LKMAATFIGNSTAIQELFKRISEQFTAMFRRKAFLHWYTGEGMDEMEFTEAESNMNDLVS   420
TBB3_HUMAN/1-450      LKMSSTFIGNSTAIQELFKRISEQFTAMFRRKAFLHWYTGEGMDEMEFTEAESNMNDLVS   420
TBB4B_HUMAN/1-445     LKMSATFIGNSTAIQELFKRISEQFTAMFRRKAFLHWYTGEGMDEMEFTEAESNMNDLVS   420
TBB5_HUMAN/1-444      LKMAVTFIGNSTAIQELFKRISEQFTAMFRRKAFLHWYTGEGMDEMEFTEAESNMNDLVS   420
TBB8_HUMAN/1-444      LKMSATFIGNNTAIQELFKRVSEQFTAMFRRKAFLHWYTGEGMDEMEFTEAESNMNDLVS   420
                      *.*: *****.*****:*:*:**:*:***:****:****.****  ** ***.*::****
```

```
TBB1_HUMAN/1-451       EYQQFQDAKAVLEEDEEVTEEAEMEPEDKGH 451
TBB2A_HUMAN/1-445      EYQQYQDATADEQGEFEEEEGEDEA------ 445
TBB2B_HUMAN/1-445      EYQQYQDATADEQGEFEEEEGEDEA------ 445
TBB4A_HUMAN/1-444      EYQQYQDATAEEGEFEEEAEEEVA------- 444
TBB3_HUMAN/1-450       EYQQYQDATAEEEGEMYEDDEEESEAQGPK- 450
TBB4B_HUMAN/1-445      EYQQYQDATAEEEGEFEEEAEEEVA------ 445
TBB5_HUMAN/1-444       EYQQYQDATAEEEEDFGEEAEEEA------- 444
TBB8_HUMAN/1-444       EYQQYQDATAEEEEDEEYAEEEVA------- 444
                       ****:***.*
```
